# THE ESSENTIAL KINASE TGGSK REGULATES CENTROSOME DIVISION AND ENDODYOGENY IN *TOXOPLASMA GONDII*

**DOI:** 10.1101/2024.09.27.615374

**Authors:** Amanda Krueger, Sofia Horjales, Chunlin Yang, William J. Blakely, Maria E. Francia, Gustavo Arrizabalaga

**Affiliations:** Department of Pharmacology and Toxicology, Indiana University School of Medicine; Laboratory of Apicomplexan Biology, Institut Pasteur de Montevideo; Department of Microbiology and Immunology, Jacobs School of Medicine and Biomedical Sciences, University at Buffalo

## Abstract

Intracellular replication is crucial for the success of apicomplexan parasites, including *Toxoplasma gondii*. Therefore, essential players in parasite replication present potential targets for drug development. In this study, we have characterized TgGSK, a glycogen synthase kinase homolog that plays an important role in *Toxoplasma* endodyogeny. We have shown that TgGSK has a dynamic localization that is concurrent with the cell cycle. In non-dividing parasites, this kinase is highly concentrated in the nucleus. However, during division, TgGSK displays a cytosolic localization, with concentration foci at the centrosomes, a key organelle involved in parasite division, and the basal end. Conditional knockdown of TgGSK determined that it is essential for the completion of the lytic cycle and proper parasite division. Parasites lacking endogenous protein levels of TgGSK exhibited defects in division synchronicity and the segregation of the nucleus and apicoplast into forming daughter cells. These phenotypes are associated with defects in centrosome duplication and fission. Global phosphoproteomic analysis determined TgGSK-dependent phosphorylation of RNA-processing, basal end, and centrosome proteins. Consistent with the putative regulation of RNA-processing proteins, global transcriptomic analysis suggests that TgGSK is needed for proper splicing. Finally, we show that TgGSK interacts with GCN5b, a well-characterized acetyltransferase with roles in transcriptional control. Conversely, GCN5b chemical inhibition results in specific degradation of TgGSK. Thus, these studies reveal the involvement of TgGSK in various crucial processes, including endodyogeny and splicing, and identify acetylation as a possible mechanism by which this essential kinase is regulated.

**AUTHOR SUMMARY:** *Toxoplasma gondii* infects nearly a third of the world’s human population. While infection is largely asymptomatic in healthy adults, in immunocompromised or immunosuppressed individuals it can lead to brain lesions and even death. Similarly, toxoplasmosis can result in stillbirth, birth defects, and blindness of a developing fetus in the case of a congenital infection. With minimal treatments for Toxoplasmosis available, it is crucial to study parasite-specific processes that could be potential drug targets for the treatment of Toxoplasmosis. In this study, we investigated the protein TgGSK that is essential for parasite survival and proper division. We showed that TgGSK may perform its essential functions through interaction with the centrosome, an organelle that plays a major role in cell division in many organisms. We also show in this study a role for TgGSK in proper processing of messenger RNAs. Taken together, we have performed an in-depth study of the functional role of the essential protein TgGSK in *Toxoplasma gondii*. Importantly, TgGSK was shown to have more similarity to plant proteins than mammalian proteins which may allow for the possibility of targeting of this protein for therapeutic treatment of toxoplasmosis.

## INTRODUCTION

*Toxoplasma gondii* is an obligate intracellular parasite that infects approximately a third of the world’s human population [1]. The parasite’s ability to be highly symptomatic in those immunocompromised or infected congenitally makes *Toxoplasma* a globally relevant pathogen [2,3]. Many of the negative effects of an uncontrolled *Toxoplasma* infection are a consequence of the lytic cycle of the highly replicative tachyzoite form, which is responsible for the acute stage of the disease. Tachyzoites actively invade any nucleated cell forming a parasitophorous vacuole in the process. After several rounds of asexual division within this vacuole, the parasite actively exits, lysing the parasitophorous vacuole and host cell in the process. Once outside the cell, the parasite quickly invades neighboring cells, allowing for propagation and continuation of the acute stage of the infection.

Intracellular division of *Toxoplasma* occurs by the divergent process of endodyogeny, through which two daughter parasites gradually form within a mature mother [4,5]. The assembly of daughter cells is supported by the inner membrane complex (IMC), which consists of a series of flattened membrane vesicles stitched together and that, along with the parasite’s plasmalemma, forms the so-called pellicle. Underneath the pellicle and serving a role in parasite shape and polarity are a set of 22 subpellicular microtubules. Early in endodyogeny, IMCs for each of the daughter cells emanate within the mother cell. Another early event in parasite division is the duplication of the centrosome, which, in addition to serving conserved roles across eukaryotes in nuclear mitosis, acts as an organizing center for subpellicular microtubules as well as serving as a contact site for various other dividing organelles in *Toxoplasma* [6]. As the IMCs of the two daughter cells grow, some organelles are made *de novo*, while the nucleus, the mitochondrion, and the plastid-like apicoplast divide between them. The IMC of the mother eventually disappears as the two nascent parasites occupy the bulk of the mother parasite. Finally, the original plasmalemma envelopes the two new cells, and a cleavage furrow extends between the daughters. The various steps that make up endodyogeny have been well described. Nonetheless, the signals and regulatory proteins controlling endodyogeny and the stepwise structural assembly of the cytoskeletal elements are not well understood.

Recently, we characterized the plant-like phosphatase PPKL, which regulates daughter cell formation in *Toxoplasma* [7]. PPKL is essential for parasite propagation, and lack of PPKL uncouples DNA duplication, which occurs normally in PPKL knockdown parasites, from daughter cell formation [7]. Knockdown of PPKL also affects the rigidity and organization of cortical microtubules, although it does not affect centrosome duplication. Interestingly, the phosphorylation of the known cell cycle regulator CRK1 is altered in PPKL knockdown parasites, suggesting that PPKL regulates parasite division by impacting the CRK-1-dependent signaling pathway [7]. PPKL is homologous to the *Arabidopsis* phosphatase BSU1, which is at the center of the plant brassinosteroid pathway [8,9]. In the absence of brassinosteroid, the phosphorylated kinase BIN2 inactivates transcription factors that regulate the expression of genes involved in various processes, including plant tissue growth, development, and stress responses. Brassinosteroid activates a cascade that results in the activation of BSU1, which in turn dephosphorylates BIN2 at a conserved tyrosine, inactivating it and leading to the transcription of brassinosteroid response genes. *Toxoplasma* does not produce brassinosteroid, and a search for other members of this well-characterized plant signaling pathway did not reveal clear homologs except for TGGT1_265330, which shows strong similarity to BIN2. As BIN2 from plants are members of the family of glycogen synthase kinase 3 (GSK3) serine/threonine kinases, we refer to TGGT1_265330 as TgGSK. A previous study, which referred to this protein as TPK3, indicated that it bears 54% homology to GSKs over the catalytic domain and showed that the recombinant protein has kinase activity and can autophosphorylate [10]. Nonetheless, the localization or function of this *Toxoplasma* kinase was not investigated.

Here, we report an in-depth analysis of TgGSK. We show that TgGSK is an essential kinase in *Toxoplasma* that displays a varying localization dependent on parasite division. Knockdown of TgGSK causes abnormal division phenotypes, including defects in the centrosome, apicoplast, and nuclear segregation. Furthermore, phosphoproteome and transcriptome analyses suggest a role in splicing for TgGSK. Finally, we show that TgGSK forms a complex with the acetyltransferase GCN5b and that inhibition of GCN5b acetyltransferase activity leads to TgGSK degradation. In sum, our work characterizes an essential kinase that regulates a variety of critical functions in the human pathogen *Toxoplasma gondii,* uncovering a potential target for therapeutic intervention.

## RESULTS

### Toxoplasma GSK is related to plant kinases

Previously, we characterized the Kelch domain-containing protein phosphatase PPKL in *Toxoplasma gondii* [7]. *Toxoplasma* PPKL is a homolog of the plant phosphatase BSU1, which is central to the brassinosteroid signaling pathway [8,9]. *Toxoplasma* does not produce brassinosteroid, and a search for other members of this well-characterized plant signaling pathway did not reveal clear homologs except for TGGT1_265330, which shows 53% identity to the PPKL substrate BIN2. BIN2 from plants are members of the family of glycogen synthase kinase 3 (GSK3) serine/threonine kinases. Thus, though this kinase had been previously referred to as TPK3 (*Toxoplasma* Protein Kinase 3), we here rename the product of TGGT1_265330 as TgGSK to better reflect its evolutionary origin and putative function. This protein is a 394 amino acid conventional kinase with an ATP binding site and an active site and shows homology to members of the GSK family of kinases. GSKs are unique among eukaryotic protein kinases (ePKs) in that they have a cluster of conserved amino acids that might perform a regulatory role. In mammalian GSK3β, these correspond to residues Q89, R92, F93, K94, and N95 [11]. TgGSK contains this cluster between amino acids 80 and 87. Moreover, existing phosphoproteomic data [12] shows that TgGSK is phosphorylated at four serine/threonine sites (T38, S208, S210, and S270) and at one tyrosine (Y211), which is conserved throughout the GSK family (**Fig 1A**). The amino acids surrounding the conserved tyrosine site are conserved across all GSKs, including in Apicomplexa, plants, and mammals (**Fig 1B**). A clustal omega alignment of TgGSK along with GSK homologs from the malarial parasite *Plasmodium*, humans, and *Arabidopsis*, shows that TgGSK and *Plasmodium* GSK cluster with the *Arabidopsis thaliana* BIN2 (**Fig. 1C**). Thus, it appears that TgGSK has greater homology to plant BIN2 than to mammalian GSK3.

**Figure 1.**
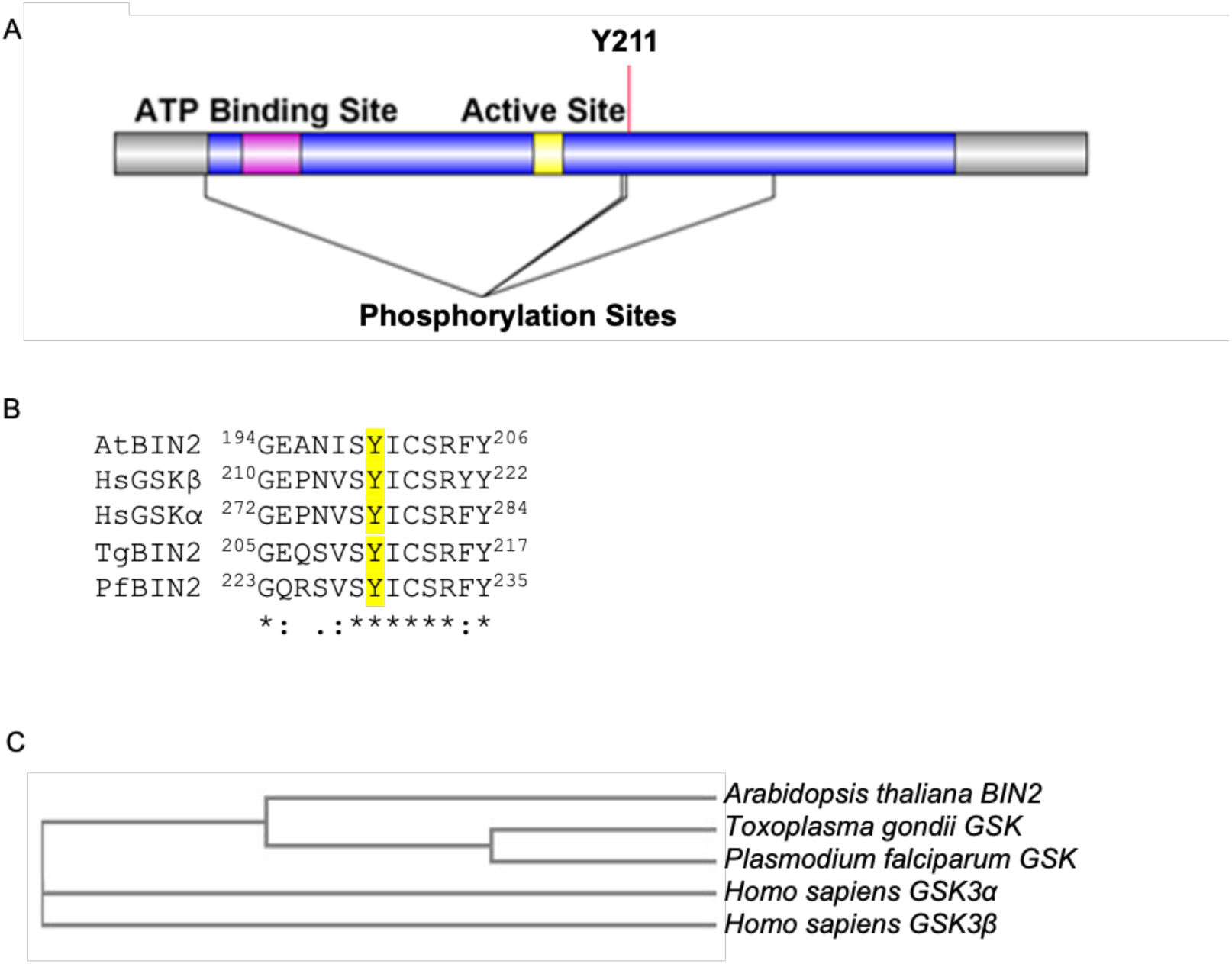
TgGSK is closely related to plant BIN2. A. Graphical representation of the TgGSK protein, with the relative position of the catalytic domain (blue), ATP binding site (magenta), and active site (yellow). Conserved acetylation (K76) and phosphorylation (Y211) sites are shown. B. Alignment of regions containing the conserved lysine and tyrosine from GSKs from *Arabidopsis thaliana* (Q39011), *Homo sapiens* (P49841, P49840), *Toxoplasma gondii* (EPT27729), and *Plasmodium falciparum* (XP_001351197) performed by Clustal Omega. C. Phylogenetic analysis of GSKs derived from Clustal Omega alignment.

### TgGSK has a dynamic localization that is cell cycle dependent

To determine the localization of TgGSK within the parasite, we used CRISPR/Cas9 to generate a strain in which the endogenous protein includes a triple hemagglutinin (3xHA) epitope tag in its carboxy terminus. Western blot analysis of the resulting strain, *ΔKu80*:TgGSK.3xHA (from here on referred to as GSK.3xHA), shows a protein of approximately 44 kDa, which matches the expected molecular weight of TgGSK (**Fig. 2A**). Using this validated strain, we performed immunofluorescence assays (IFA) of intracellular parasites probing for HA to observe TgGSK and for IMC3 to monitor the inner membrane complex (IMC) (**Fig. 2B**). IMC3 is present in both mother and daughter parasites, which allows us to differentiate between dividing (those with daughter parasites within) and non-dividing (those without daughter parasites) parasites. Interestingly, we noted that while in non-dividing parasites, TgGSK is more concentrated in the nucleus, in those undergoing division TgGSK is evenly distributed throughout the entire cell (**Fig. 2B**). To explore this dynamic localization, we monitored TgGSK’s localization in parasites at various stages of division (**Fig. 2C**). Throughout all stages of division, we saw TgGSK distributed throughout the cell, while in non-dividing parasites TgGSK consistently localized to the nucleus. While the difference in GSK localization in dividing and non-dividing parasites is evident, we aimed to quantify this observation. For this purpose, we imaged 20 non-dividing parasites and 20 parasites in the late division stage and used ImageJ to quantify the fluorescent intensity of the HA signal in the cytosol and the nucleus. As expected, this analysis determined a higher ratio of nuclear over cytosolic signal in non-dividing parasites (**Fig. 2D**).

**Figure 2.**
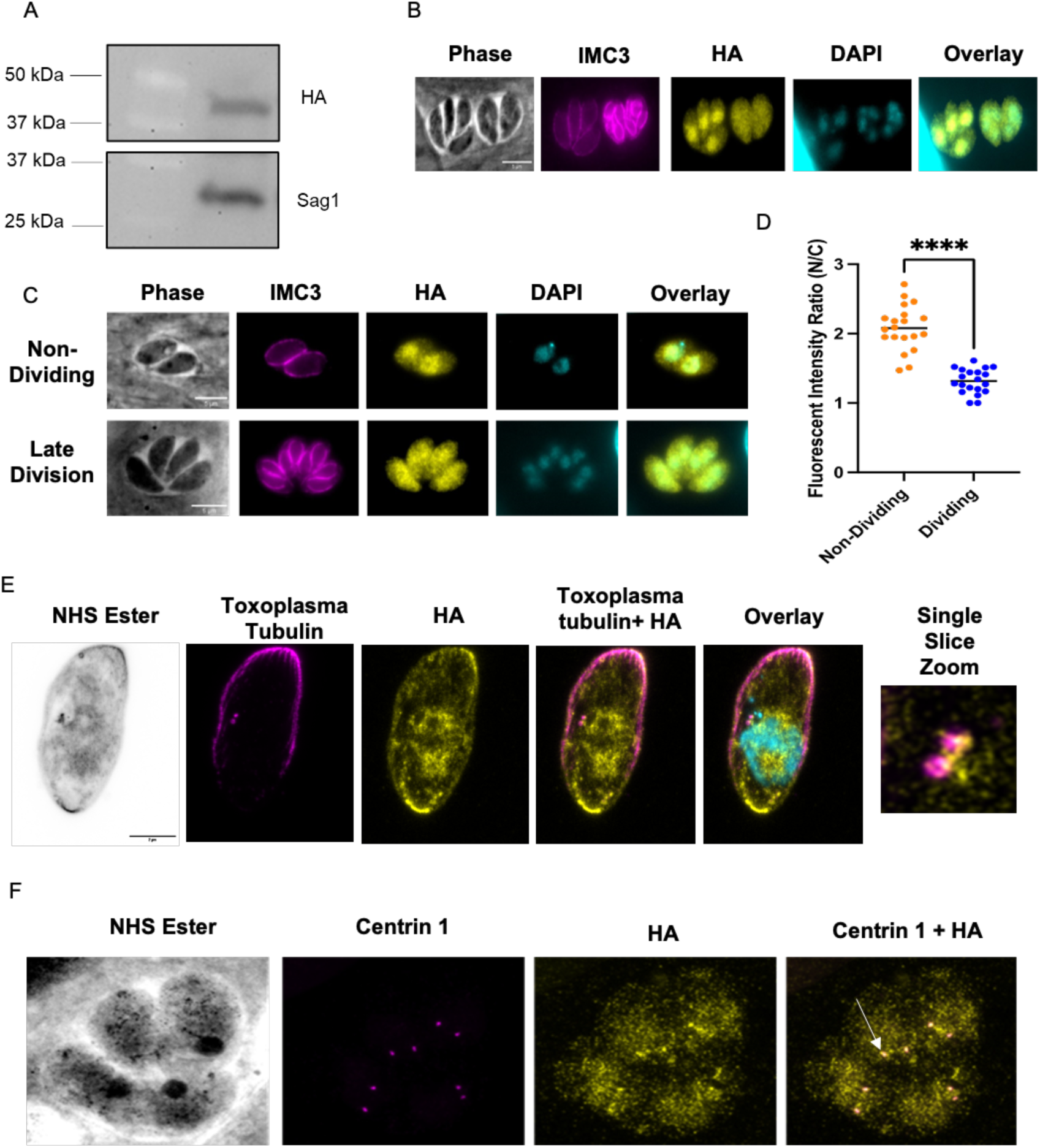
TgGSK has a dynamic, cell cycle-dependent localization. A. Western blot of protein extract from parasites in which the endogenous TgGSK includes a 3xHA epitope tag. Blots were stained for HA (top) and Sag 1 (bottom). B and C. Parasites of the TgGSK.3xHA strain were grown intracellularly for 24 hours before performing IFA using antibodies against HA and IMC3, and the DNA stain DAPI. In C, parasites in the top panels are not dividing, while those in the bottom panel are undergoing division. D. Quantification of the fluorescent intensity ratio of nucleus to cytosol in non-dividing and dividing parasites; n=20 per condition, p<0.0005. E. Expansion microscopy of a non-dividing parasite staining for *Toxoplasma* tubulin (magenta), HA (yellow), and DRAQ5 (cyan, color in overlay) to visualize parasite structure and centrosomes, TgGSK, and nuclear material, respectively. A single slice zoomed image of the centrosomes is shown to closer visualize TgGSK localization to this organelle. F. Expansion microscopy of four dividing parasites in a vacuole staining for Centrin 1 and HA to visualize TgGSK’s colocalization with the centrosomes. Arrowhead shows an example of TgGSK colocalized with a centrosome.

To obtain a more detailed understanding of TgGSK’s localization, we performed ultrastructure expansion microscopy (UExM), which allows for higher-resolution analysis. As with standard IFA, we detect TgGSK within the nucleus. Interestingly, we also detect TgGSK in areas of tubulin concentration, which are reminiscent of centrosomes, as well as in the basal end of the parasites (**Fig. 2E**). Given the apparent localization of TgGSK to the centrosome, we co-stained for TgGSK and Centrin 1. Indeed, TgGSK and Centrin 1 appear to localize to the same area, confirming TgGSK’s concentration around the centrosomes in both non-dividing and dividing parasites (**Fig. 2F**). In sum, IFA and UExM analyses show that TgGSK has a dynamic localization dependent on the division stage and is present at the centrosomes, suggesting that TgGSK may play a role in the regulation of parasite replication.

### TgGSK IS ESSENTIAL FOR PARASITE SURVIVAL

In a *Toxoplasma* genome-wide CRISPR screen, TgGSK was assigned a fitness value of -4.12 [13], which suggests that this protein is essential and, therefore, a full knockout would likely not be possible. Accordingly, we used a conditional knockdown approach to investigate the function of TgGSK. For this purpose, we replaced the TgGSK promoter with the tetracycline regulatable promoter (TATi) [14] in the GSK.3xHA strain using CRISPR/Cas9 (**Fig. 3A**). IFA showed that TgGSK’s localization is unchanged in the resulting strain (TATi-GSK.3xHA) (**Fig. 3B**). Importantly, western blot analysis shows that TgGSK expression is significantly reduced after 42 hours of treatment with the tetracycline analog aTC, with near complete lack of protein at 72 hours (**Fig. 3C**). To determine if TgGSK is required for parasite propagation, we performed a plaque assay with the parental and the TATi-GSK.3xHA strains with and without aTC. Consistent with the results of the CRISPR screen, TATi-GSK.3xHA parasites grown in the presence of aTC failed to form plaques (**Fig. 3D and E**), indicating that TgGSK is essential for parasite survival and propagation.

**Figure 3.**
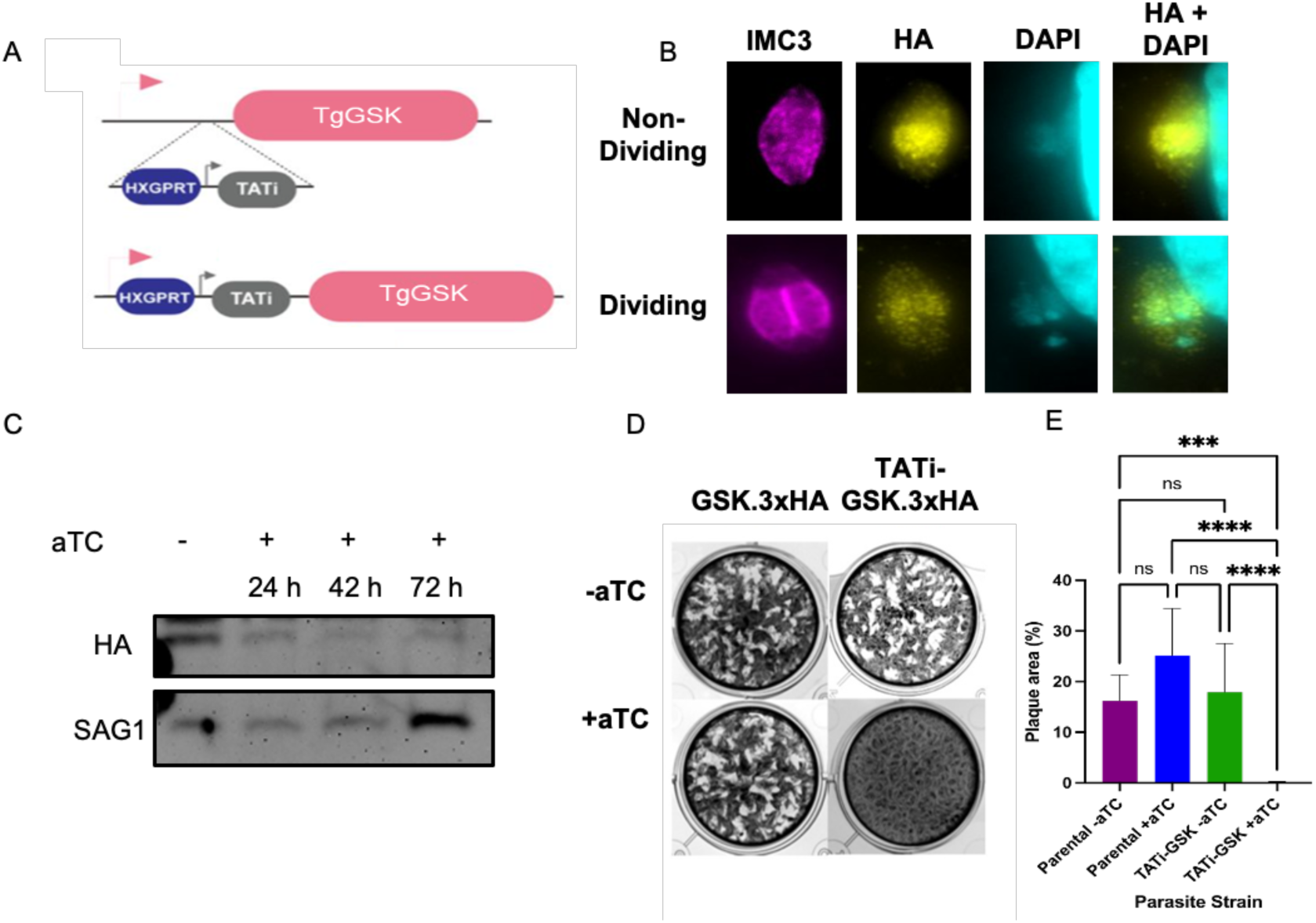
TgGSK is essential for propagation in tissue culture. A. Graphical representation of the strategy used to engineer a TgGSK conditional knockdown strain using the TATi tetracycline regulatable system. B. Parasites of the TATi-GSK.3xHA strain were grown in culture for 24 hours to perform IFA with antibodies against IMC3 and HA and the DNA stain DAPI without the addition of aTC. C. Western blot of protein extract from TATi-GSK.3xHA parasites grown in no aTC (-) or in aTC for 24, 42, or 72 hours. Blots were probed for HA to monitor TgGSK or Sag1 as loading control. D. Plaque assay of parental (GSK.3xHA) and TATi-GSK.3xHA knockdown parasites incubated for 6 days with or without aTC E. Quantification of plaque assay. ****: p<0.0005, ***: p<0.005, ns: no significance, n=9 wells per condition (3 biological replicates with 3 experimental replicates each).

### KNOCKDOWN OF TgGSK CAUSES ABNORMAL DIVISION PHENOTYPES

To explore the impact of TgGSK’s loss on cell division, we performed IFAs on the TATi-GSK.3xHA strain grown with and without aTC. Staining for IMC3 reveals significant division defects after 42 hours of aTC treatment, at which time point there is still some TgGSK present, albeit at reduced levels (**Fig. 4A**). The most common phenotypes observed include asynchronous division, incomplete nuclear segregation, and vacuoles with parasites of abnormal shape (**Fig. 4A**). Quantification of 100 vacuoles over three experimental replicates showed that, while 76.5%±2.2% of parasites appear to divide normally in the absence of aTC, only 22.9%±3.18% of parasites exhibit normal division in the presence of aTC (**Fig. 4B**). Among the vacuoles grown with aTC that show aberrant division, 51.9%±1.5% exhibit asynchronous division, 28.6%±2.4% uneven segregation, and 19.4%±2.9% abnormally shaped parasites (**Fig. 4C**). When parasites were allowed to grow in aTC for 96 hours, there was an exacerbation of all phenotypes as the parasites appear to continue dividing unsuccessfully (**Fig. 4D**). Notably, significant defects are seen in parasite structure, with acetylated tubulin not being organized adequately into individual fully closed parasites, and in nuclear segregation. Interestingly, there also seems to be a defect in chromatin condensation, as shown by diffuse staining for the histone marker H2 (**Fig. 4D**). Overall, analysis of the TgGSK knockdown strain reveals a function for this kinase in parasite division.

**Figure 4.**
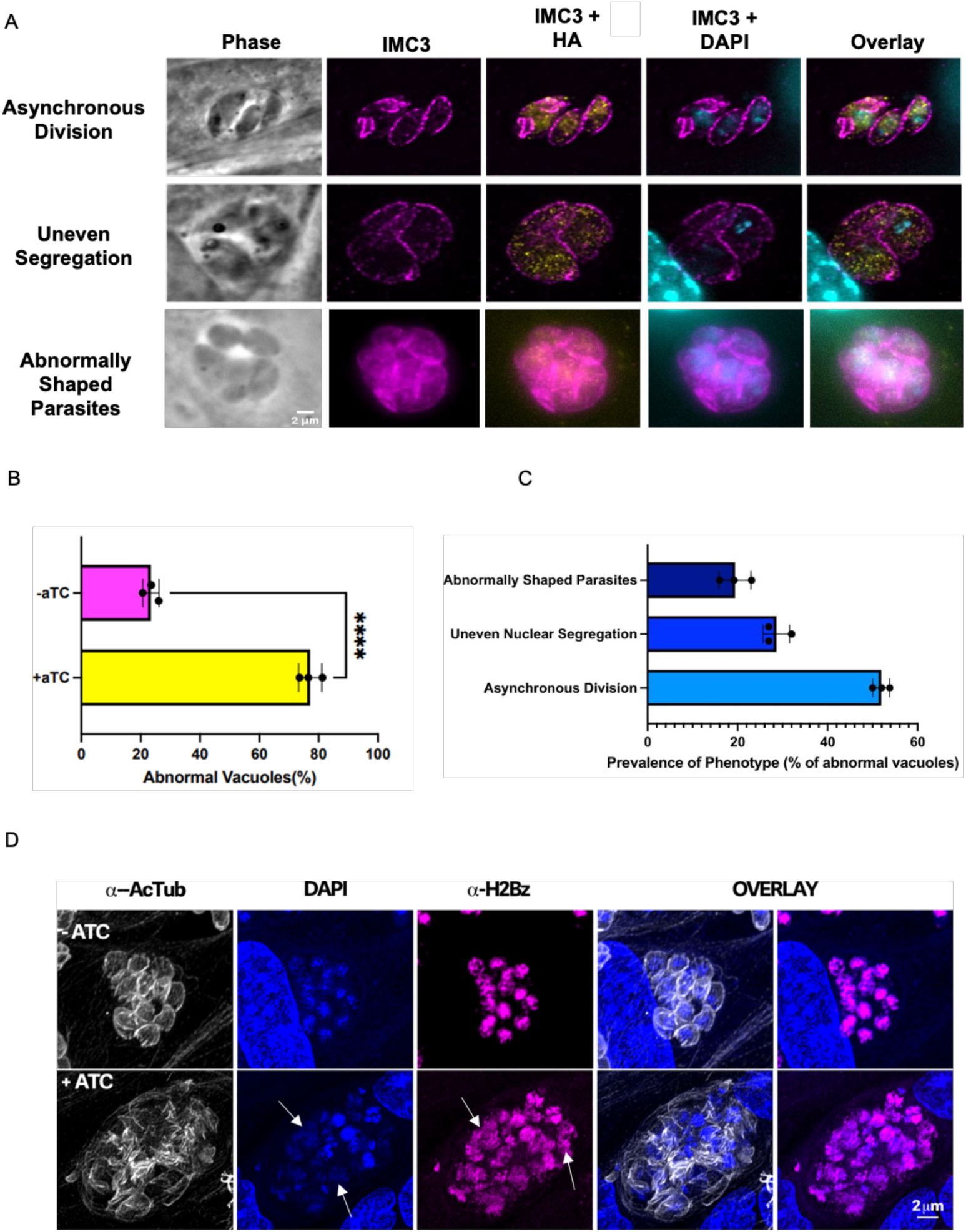
Knockdown of TgGSK causes abnormal division phenotypes. A. IFA of TgGSK knockdown parasites after 42 hours of incubation with aTC. IMC3, HA, and DAPI staining visualize mother and daughter cell IMC, TgGSK, and nuclear material, respectively. Representatives of the main phenotypes observed are shown B. Quantification of the percentage of normal vacuoles in TATi-GSK parasites with and without 42-hour incubation with aTC. N= 3, 100 total parasites over the replicates. The error percentage shown is the standard deviation. C. Quantification of the different division phenotypes seen after 42 hours of TgGSK knockdown. D. IFA showing division phenotypes of TgGSK knockdown parasites after 96 hours of incubation with aTC. Acetylated tubulin, DAPI, and H2Bz staining visualize parasite structure, nuclear material, and histones, respectively. Arrowheads point to diffuse nuclear material and H2 histone.

### TgGSK KNOCKDOWN CAUSES CENTROSOME ABNORMALITIES

Since we detected TgGSK in the centrosomes and observed nuclear segregation defects in TgGSK knockdown parasites, we asked if knockdown also caused centrosome abnormalities. Indeed, using IFA staining for Centrin 1 and the mitotic spindle marker EB1, we detect abnormalities in centrosome morphology upon TgGSK knockdown (**Fig. 5A**). Specifically, we observe elongated centrosomes and some that seem unable to undergo fission. We quantified the distribution of EB1 and centrin per parasite nucleus from our IFA images using ImageJ. In a normal parasite culture, where around 30% of parasites are dividing, the average amount of centrosomes per nucleus should be around 1.25, with non-dividing parasites having one centrosome and dividing parasites having two [13]. EB1 recruitment to the mitotic spindle accompanies centrosome duplication and is only detectable in dividing parasites. While there was no significant difference in the number of nuclei displaying EB1 foci between control and TgGSK knockdown parasites, there was a significant difference in the number of centrosomes associated with each parasite nucleus (**Fig. 5B**). The aTC-treated parasites had a lower average number of centrosomes per nucleus, with some vacuoles displaying only one centrosome for four parasite nuclei (**Fig. 5B**). Importantly, UExM confirmed the various phenotypes observed by IFA (e.g. abnormal parasite structure and nuclear segregation defects) and highlighted the abnormally shaped centrosomes at a higher resolution (**Fig. 5C**). Given the centrosome’s key role in parasite division and organellar segregation, the various division-related phenotypes observed in the TgGSK knockdown parasites might be a consequence of the centrosome segregation defects present in the mutant.

**Figure 5.**
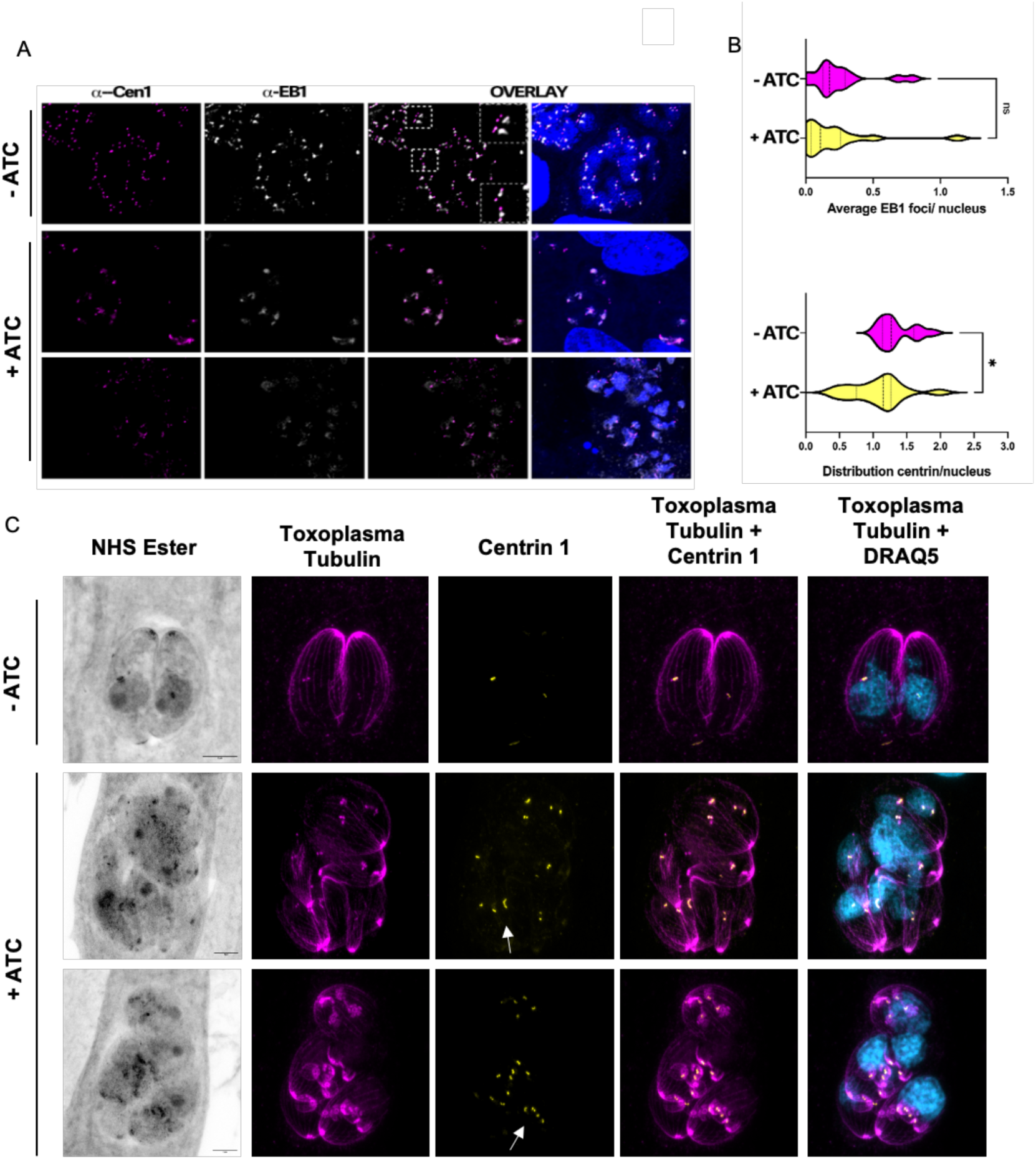
TgGSK knockdown parasites have a centrosome duplication defect. A. IFA TATi-GSK parasites treated with aTC for 96 hours stained for centrin 1 (centrosomes) and EB1 mitotic spindles). Outlined boxes show parasites with abnormally dividing centrosomes. B. Quantification of centrin and EB1 signal in (A) measuring distribution of signal per parasite nucleus. N= 3 replicates with 100 vacuoles quantified. Statistical analysis: two-tailed t-test with Welsh’s correction. *: p<0.05 ns: no significance. C. Expansion microscopy of TATi-GSK parasites treated with aTC for 42 hours. Arrows indicate elongated centrosomes that seem unable to undergo fission.

### KNOCKDOWN OF TgGSK AFFECTS APICOPLAST DIVISION

It is known that the centrosomes also coordinate the segregation of other organelles, including the apicoplast [15]. Accordingly, we monitored apicoplast division and segregation by monitoring the apicoplast marker CPN60 (**Fig. 6A**). While parasites grown in the absence of aTC averaged around one apicoplast per parasite nucleus, aTC treated parasites averaged one apicoplast for every four parasite nuclei (**Fig. 6B**). Therefore, it appears that, in addition to abnormal nuclear division and segregation, apicoplast division and segregation are also disrupted in TgGSK knockdown parasites.

**Figure 6.**
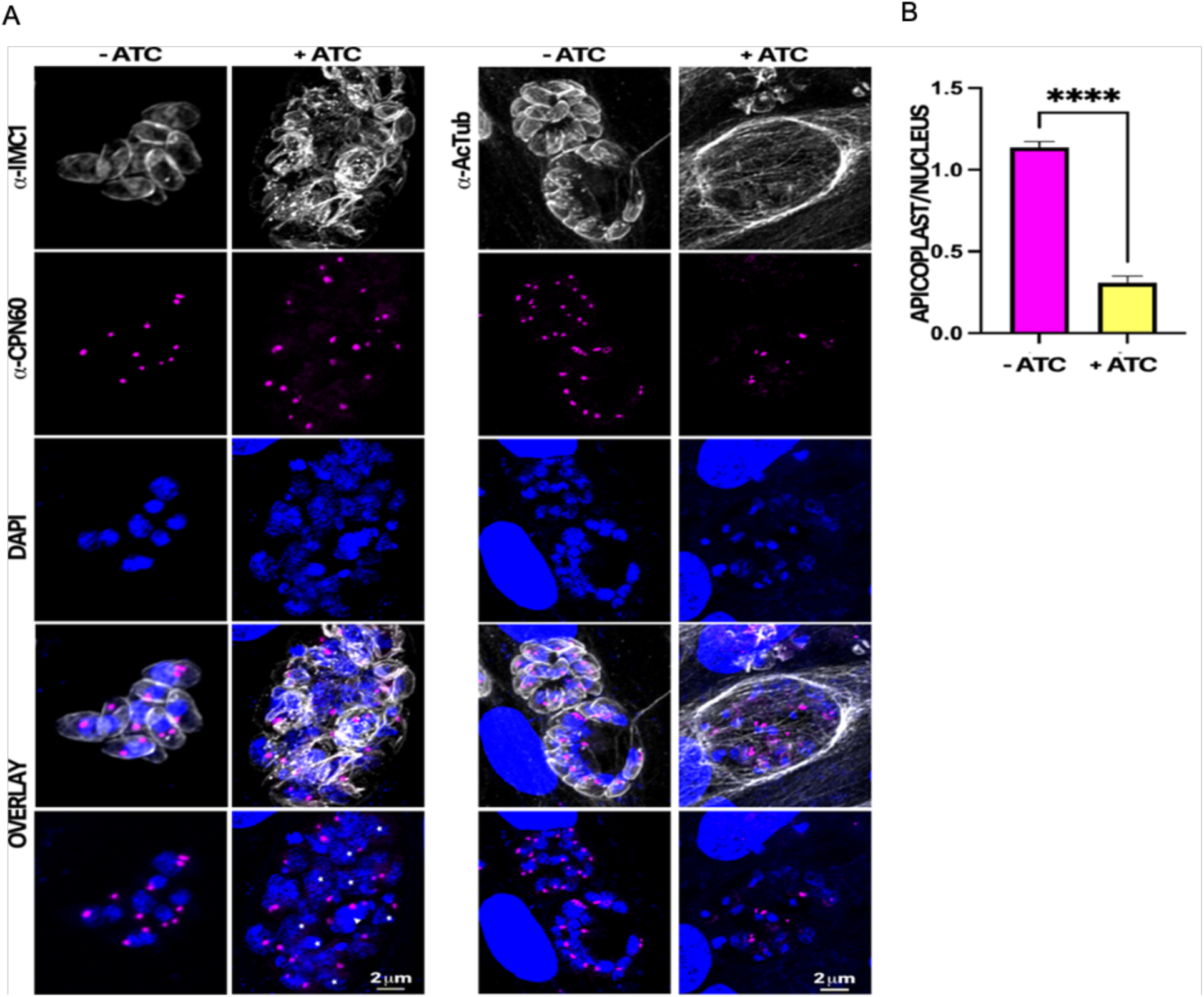
TgGSK knockdown parasites display a reduction in apicoplast material. A. IFA of TATi-GSK parasites after 96 hours of aTC treatment stained for the apicoplast marker CPN60. IMC1 and DAPI staining are also used to visualize parasite structure and nuclear material. B. Quantification of the number of apicoplast foci per parasite nucleus in TgGSK knockdown versus control parasites. ****: p<0.0005 n=3 experimental replicates, 100 total parasites quantified.

### CENTROSOMAL, BASAL END, AND SPLICING PROTEINS SHOW TgGSK-DEPENDENT PHOSPHORYLATION

Since TgGSK has the structure of a conventional kinase and is a member of the GSK family, we performed global phosphoproteome analysis to determine proteins that have TgGSK-dependent phosphorylation. We found 27 proteins with peptides that had decreased phosphorylation and 40 proteins with peptides that had increased phosphorylation in TgGSK knockdown parasites compared to parental (Log2FC>0.5, p<0.05) (**Fig. 7A and supplemental dataset 1).** Using ToxoDB and StringDB, we identified a few enriched pathways involving some of the 67 proteins displaying TgGSK-dependent phosphorylation. Three of the 27 proteins that had decreased phosphorylation in the knockdown were RNA-binding or known splicing proteins (**Fig. 7B**). Another three proteins with decreased phosphorylation were members of the MyoC glideosome complex in the basal end of the parasite (**Fig. 7B**). Interestingly, of these six proteins of interest, five of them were differentially phosphorylated at an S/TXXXS/T motif, typical of GSK substrates [16]. In addition to splicing and basal end proteins, the protein that had the highest increase in phosphorylation in the knockdown was Centrin 2 (**Fig. 7B**). In addition, analysis of the hypothetical proteins with TgGSK-dependent phosphorylation revealed that 18% of those could be involved in RNA metabolism **(Supplemental figure 1).** Overall, phosphoproteome analysis suggests that TgGSK might influence the regulation of proteins related to the centrosomes, basal complex, and splicing.

**Figure 7.**
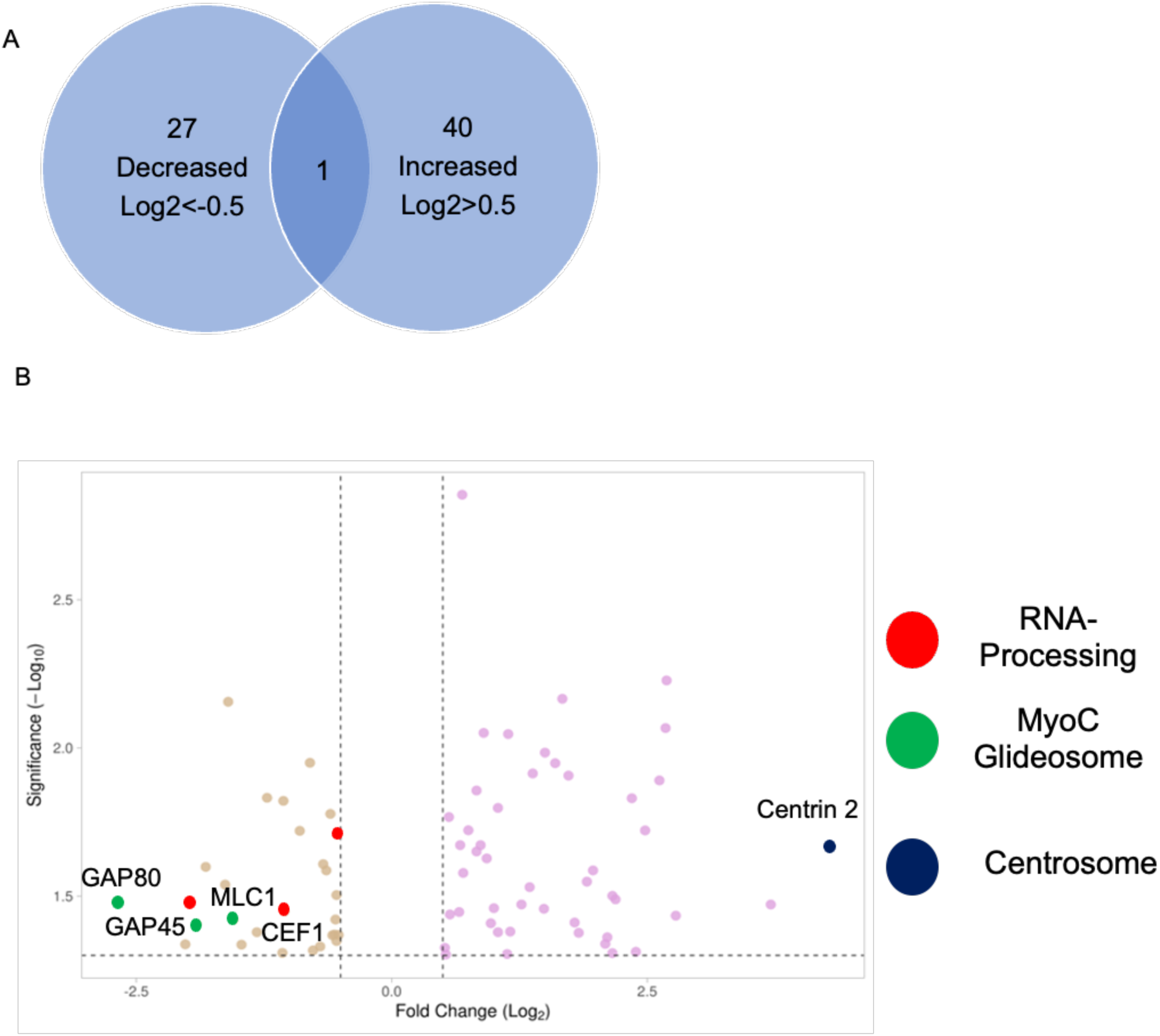
Global Phosphoproteome analysis reveals TgGSK-dependent phosphorylation events. A. Number of proteins that had increased or decreased phosphorylation after 24 hours of TgGSK knockdown. One protein had peptides with both increased and decreased phosphorylation. B. Volcano plot of all differentially phosphorylated proteins with a log2fold change >0.5 and p<0.05. Splicing, myoC glideosome, and centrosome proteins are highlighted.

### TgGSK INTERACTS AND IS REGULATED BY THE GCN5B COMPLEX

To further understand the role of TgGSK and the phenotypes associated with its loss, we performed immunoprecipitation followed by mass spectrometry to identify putative interacting partners. Interestingly, the top nine significant TgGSK interacting proteins were all in the nucleus, with eight of them being members of the GCN5b complex (**Table 1 and supplemental dataset 2)**. The GCN5b complex is present in the nucleus, where it acetylates histones to open chromatin for gene transcription [17].

**Table 1.**
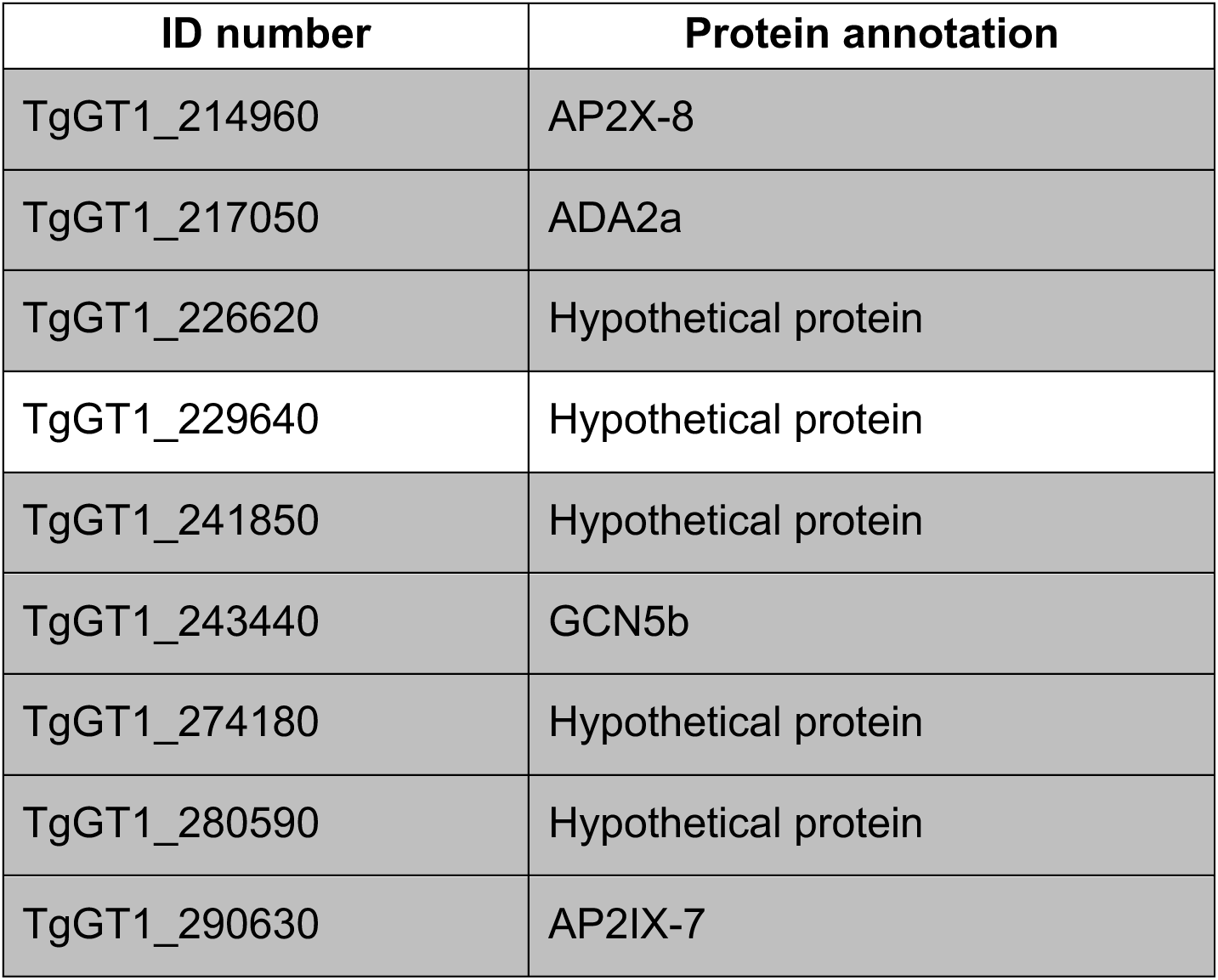
TgGSK interactors. ID number and annotation of putative interactors of TgGSK identified in both IPs performed with a ratio of experimental over control higher than 5. Members of the GCN5b complex are shown in grey.

| ID number | Protein annotation |
| --- | --- |
| TgGT1_214960 | AP2X-8 |
| TgGT1_217050 | ADA2a |
| TgGT1_226620 | Hypothetical protein |
| TgGT1_229640 | Hypothetical protein |
| TgGT1_241850 | Hypothetical protein |
| TgGT1_243440 | GCN5b |
| TgGT1_274180 | Hypothetical protein |
| TgGT1_280590 | Hypothetical protein |
| TgGT1_290630 | AP2IX-7 |

While it is plausible that TgGSK regulates members of the GCN5b complex via phosphorylation, we did not identify any of them as part of the TgGSK-dependent phosphoproteome. Accordingly, we explored whether, alternatively, the GCN5b complex might regulate TgGSK. For this purpose, we treated parasites with Garcinol, an acetyltransferase inhibitor that has been shown to specifically inhibit the histone acetylation activity of GCN5b in *Toxoplasma* [18]. We treated GSK.3xHA parasites with 0, 2, or 4µM of Garcinol overnight and monitored TgGSK localization by IFA. Interestingly, we found that while there was no difference in TgGSK localization pattern after this incubation time, the overall TgGSK signal was reduced in a Garcinol dose-dependent manner, with the signal being absent after treatment with 4µM of Garcinol (**Fig. 8A**). To confirm these results, we performed western blot analysis using the same Garcinol concentrations. We saw a reduction in TgGSK protein levels after treatment with 2µM of Garcinol and a complete absence of protein after treatment with 4µM of Garcinol (**Fig. 8B**). As a control, we also stained for aldolase, which did not seem to be affected across all Garcinol concentrations tested (**Fig. 8B**). Interestingly, previously published data showed that TgGSK expression level was not changed by Garcinol treatment, further suggesting that TgGSK is regulated by GCN5b at the protein level [18]. Therefore, it appears that TgGSK’s protein expression or stability might be regulated by GCN5b.

**Figure 8.**
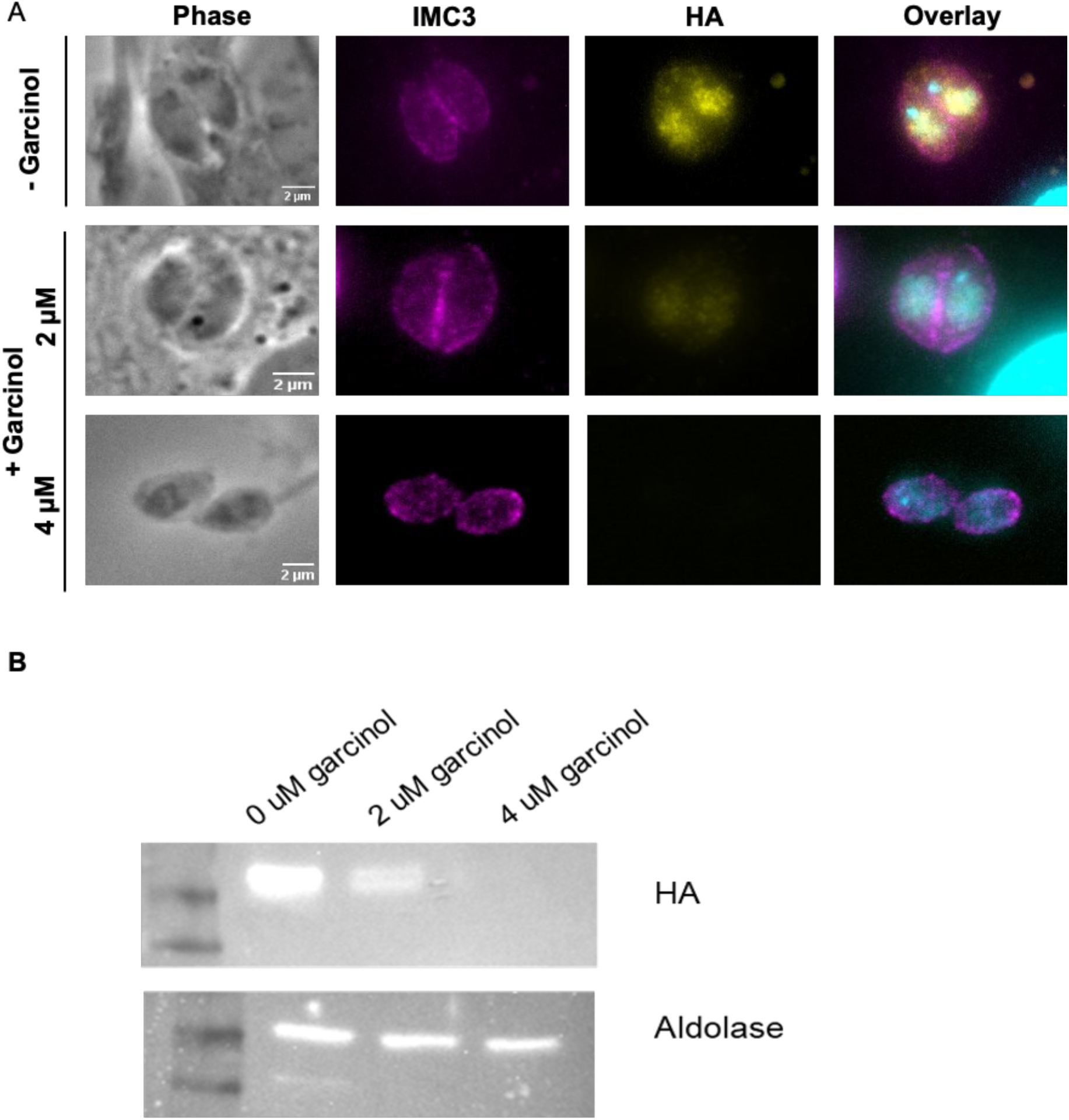
Acetylation by GCN5b stabilizes TgGSK. A. IFA of non-dividing GSK.3xHA parasites treated with 0, 2, or 4 µM Garcinol for 18 hours. Staining was done for IMC3, HA, and DAPI to visualize IMC, TgGSK, and nuclear material, respectively. B. Western blot analysis of TgGSK protein levels after 18 hours of Garcinol treatment. The cytosolic protein aldolase was used as a control.

### TgGSK KNOCKDOWN CAUSES DIFFERENTIAL TRANSCRIPTION AND SPLICING

As we detected TgGSK in the nucleus and determined that it is in a complex with transcription factors we investigated the effect of TgGSK knockdown on global transcription. For this purpose, we performed RNAseq of the TATi-GSK.3xHA strain grown for 18 hours with or without aTC. We used 18 hours, at which time point there is still TgGSK protein, to avoid transcript changes associated with parasite death. We found that there were 405 genes downregulated and 157 genes upregulated when TgGSK was knocked down (Log2FC>0.5, p<0.05) (**Fig. 9A and supplemental dataset 3)**. The differentially regulated genes were members of many different pathways, with most of them being either hypothetical or not falling into any enriched pathway (**Fig. 9B**). Comparison of the dysregulated genes with those whose promoters are known to be bound by the GCN5b complex did not reveal any enrichment for GCN5b regulated genes [17].

**Figure 9.**
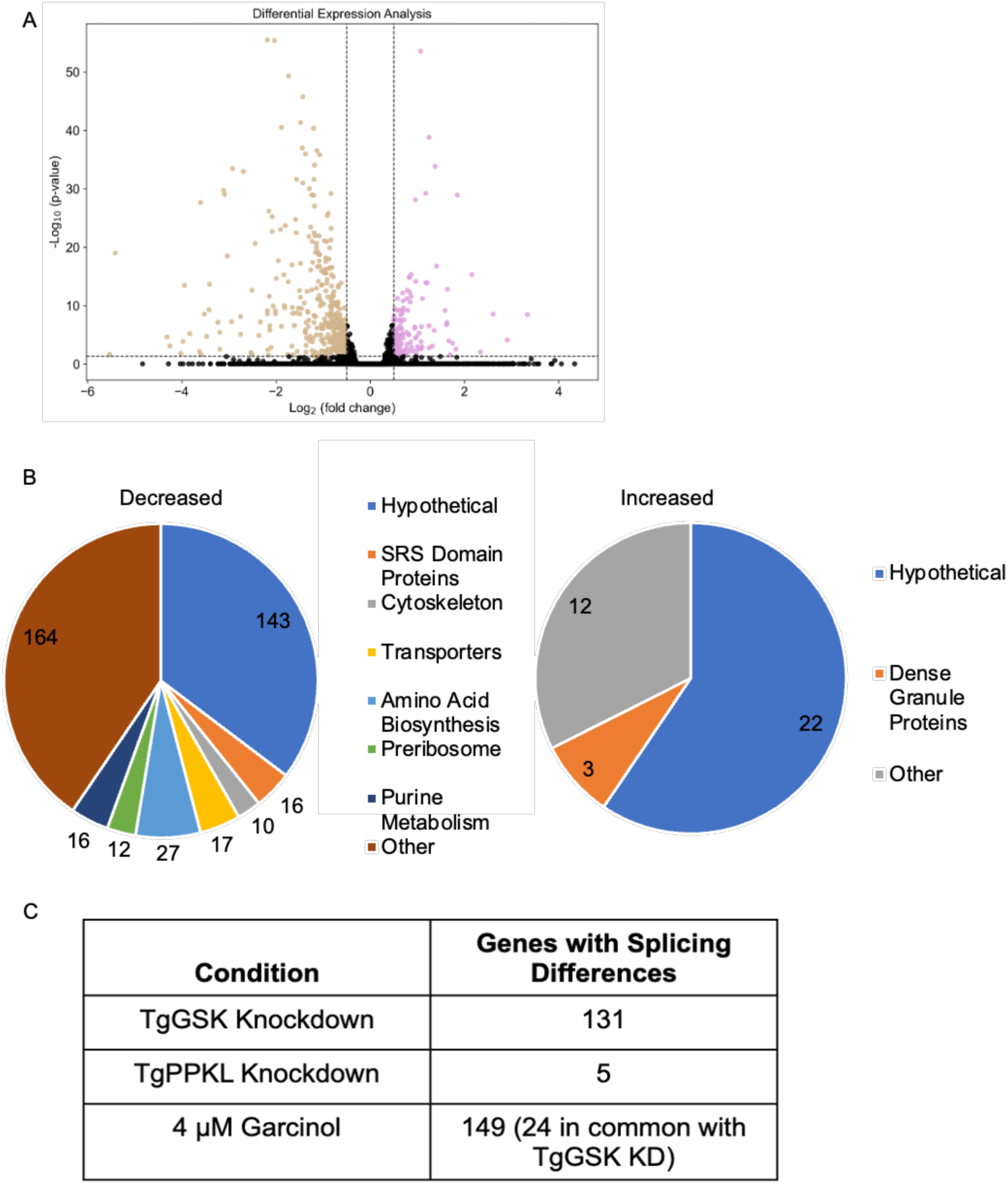
Global transcriptome analysis of TgGSK knockdown parasites. A. Volcano plot showing all differentially transcribed genes with a log2fold change >0.5 and p<0.05 after 18 hours of TgGSK knockdown. B. Biological processes with enriched transcriptome changes as identified by ToxoDB and StringDB. C. Genes that were differentially spliced after TgGSK knockdown, TgPPKL knockdown, and treatment with 4 µM Garcinol.

As we identified several RNA processing proteins that had TgGSK-dependent phosphorylation, we investigated the RNAseq data for splicing variants. Interestingly, we found that 131 genes had exon differences in TgGSK knockdown parasites compared to parental (**Fig. 9C**). Since this data was taken at 18 hours of aTC treatment, we infer that these splicing differences were not due to parasite death. However, as an additional control, we also analyzed exon differences in the transcriptome of the TgPPKL knockdown parasites, which, like TgGSK, is essential, and its disruption causes division-related defects [7]. We found that compared to parental, only five genes were differentially spliced (**Fig. 9C**). In addition, since the garcinol treatment resulted in loss of TgGSK, we mined previously published transcriptomic data from garcinol-treated parasites [18] to assess the effects on splicing. This analysis revealed 149 differentially spliced transcripts in parasites treated with 4 µM Garcinol (**Fig. 9C**). Interestingly, 24 of these overlapped with transcripts that were differentially spliced in the TgGSK knockdown (**Fig. 9C**). Overall, these data suggest that TgGSK plays a role in proper splicing and this effect is seen after both TgGSK transcriptional knockdown and protein degradation.

## DISCUSSION

Members of the glycogen synthase kinase (GSK) protein family are serine/threonine kinases present in many organisms, including mammals and plants [19]. Most mammalian species encode for two GSKs, GSK3α and GSK3β, with hundreds of substrates that play roles in cellular proliferation and migration, glucose regulation, and apoptosis [20]. By contrast, most plant species encode 10 GSKs, which fall into four major groups and are involved in plant growth, development, and stress response [21]. All GSKs have a single conserved tyrosine residue that can be phosphorylated [22]. This residue has been shown to interact in the binding pocket to control kinase regulation [23]. In Arabidopsis, this tyrosine is dephosphorylated by the protein phosphatase BSU1, leading to its inactivation [9]. Interestingly, *Toxoplasma gondii,* like other parasites of the phylum Apicomplexa, encode for two putative GSKs (TGGT1_265330 and TGGT1_266910), which are, by and large, uncharacterized. In the current study, we investigated the localization and function of TGGT1_265330, which we refer to as TgGSK. We found that TgGSK localization is dependent on the cell cycle and that depletion of TgGSK impacts parasite daughter formation, nuclear segregation, centrosome dynamics and fission, and apicoplast dynamics. Our findings also demonstrated that TgGSK might play a role in the regulation of splicing and that its stability is dependent on the GCN5b lysine acetyltransferase.

The asexual division of *Toxoplasma* occurs by the unusual process of endodyogeny, which is defined by the gradual formation of two daughter parasites within a mature one. The centrosome is an essential component of this division process, with each daughter parasite forming around a centrosome, which undergoes fission in early S phase [25]. The centrosome has been shown to nucleate spindle microtubules during mitosis [25]. Later, during cytokinesis, the centrosome organizes the scaffolding of daughter cell components to allow for nuclear fission and correct organelle segregation into daughter parasites [25]. One of the organelles that is associated with the centrosomes during division is the apicoplast, a non-photosynthetic plastid organelle [15]. Before apicoplast fission, the organelle elongates, with each end interacting with a centrosome. Therefore, the centrosomes act to correctly orient the apicoplast to allow for proper fission. By UExM, we were able to visualize TgGSK at the centrosomes. In addition, knockdown of TgGSK led to defects in centrosome number, with not every parasite nucleus having an associated centrosome. We also observed defects in centrosome duplication, with centrosomes that appeared to be duplicated but unable to undergo fission. These centrosome abnormalities could underlie the other division phenotypes we observe, such as nuclear and apicoplast segregation defects. Interestingly, we also observed TgGSK-dependent phosphorylation of Centrin 2, a protein that is localized to the centrosomes and has been shown to be essential for parasite division and correct centrosome segregation [26]. In our study, Centrin 2 phosphorylation was increased during TgGSK knockdown, suggesting that it is not a TgGSK substrate and that its regulation of Centrin 2 is indirect. Interestingly, Centrin 2 has also been shown to be localized to the basal body [26]. As we also detect TgGSK within the basal end of the parasite, it is unclear whether the functional relationship between these two proteins occurs at the centrosome or in the basal body.

A role for GSKs in centrosome regulation is also observed in other organisms. For example, inhibition of GSK3β in human cancer cells results in centrosome dysregulation and abnormal mitosis [27]. Similarly, in HeLa cells, GSK3β plays a role in the organization of microtubule arrays derived from the centrosomes [28]. The knockdown of GSK3β reduced the amount of centrosomally focused microtubules and caused the mislocalization of various centrosomal proteins [28]. Though recruitment of the mitotic spindle binding protein EB1 is not altered in the absence of TgGSK, further studies would need to be done in *Toxoplasma* to determine if centrosomal proteins or mitotic factors are negatively affected in the absence of TgGSK.

Intriguingly, we determined that the phosphorylation state of three RNA-binding proteins (TGGT1_264610, TGGT1_275480, and TGGT1_304630) are altered in the absence of TgGSK. While TGGT1_264610 and TGGT1_304630 are characterized as putative RNA binding proteins in *Toxoplasma,* StringDB analysis characterized them as proteins related to splicing. TGGT1_275480 is homologous to the pre-splicing factor CEF1. Interestingly, all three of these proteins contain the S/TXXXS/T motif, suggesting that they may be direct substrates of TgGSK [17]. As with GSKs in the centrosomes, there is evidence from other organisms highlighting the potential role GSKs play in splicing. A study in mouse embryonic stem cells found that inhibition of GSK3 altered the splicing of 188 mRNAs [29]. GSK was also shown to interact with multiple SR family splicing proteins and various other RNA-binding proteins [29]. In human T-cells, phosphorylation of the nuclear RNA biogenesis protein PSF by GSK3 controls the alternative splicing of CD45 [30]. While there is no direct evidence for a role in splicing for the plant GSK homolog BIN2, mapping of the BIN2 signaling network identified 13 RNA processing proteins [31]. Thus, it is plausible that *Toxoplasma* TgGSK directly or indirectly regulates the function of RNA processing proteins. Consistent with this idea, we observed an increase in alternatively spliced transcripts upon knockdown of TgGSK.

One of the most intriguing findings of our studies is the possible regulation of TgGSK by the lysine acetyltransferase GCN5b. We observed that TgGSK interacts with a well-characterized GCN5b-containing complex. *Toxoplasma* encodes for two GCN5 proteins, with GCN5b being essential for parasite viability [17]. The GCN5b complex is in the nucleus, where it performs its primary function of acetylating histones [17]. GCN5b has been shown to be present in a complex that includes the ADA2a adaptor protein and various plant-like AP2 transcription factors [32]. There appear to be two distinct stable GCN5b-containing complexes in *Toxoplasma*, one which includes the putative transcription factors AP2X8 and AP2IX7 and the other which includes AP2XII4 and AP2VIIa5 [32]. Our results indicate that TgGSK interacts with the complex that includes AP2X8 and AP2IX7 (**Table 1**). Interestingly, previous characterization of the *Toxoplasma* GCN5b complexes identified TgGSK as an interactor, which validates the interaction between TgGSK and this complex [32].

Not only did we detect a physical interaction between TgGSK and the GCN5b complex, but we also observed a specific loss of TgGSK upon GCN5b inhibition by Garcinol. These results bring up the possibility that GCN5b regulates TgGSK via acetylation. While acetylation of histones is canonical for histone acetyltransferases, there have been many studies showing acetylation of non-histone proteins in various organisms [33,34]. Non-histone protein acetylation has been found to play a broad diversity of roles, including protein folding and stability [35]. Consistent with a possible role of acetylation in the regulation of TgGSK, a global acetylome study in *Toxoplasma* identified lysine acetylation on TgGSK at residue K13 in extracellular parasites. Intriguingly, a whole proteome mapping of ubiquitination showed that K13 is also ubiquitinated [36]. Cross-talk between acetylation and ubiquitination of the same lysine is well known and is of particular importance in the context of protein stability, where lysine acetylation can block proteasome-mediated degradation by lysine acetylation [37]. Thus, our data points to a novel mechanism of TgGSK regulation via the competition between acetylation and ubiquitination (**Fig. 10**). We propose that TgGSK regulation could involve its acetylation within the nucleus by GCN5b at residue K13 before being trafficked to the cytosol and centrosomes in preparation for cell division. Once division has finished, TgGSK could be deacetylated and ubiquitinated at K13, causing its degradation (**Fig. 10**). Further studies focused on whether K13 in TgGSK plays a direct role in the stability and function of the protein are thus warranted as it would elucidate a novel mechanism of GSK regulation.

**Figure 10.**
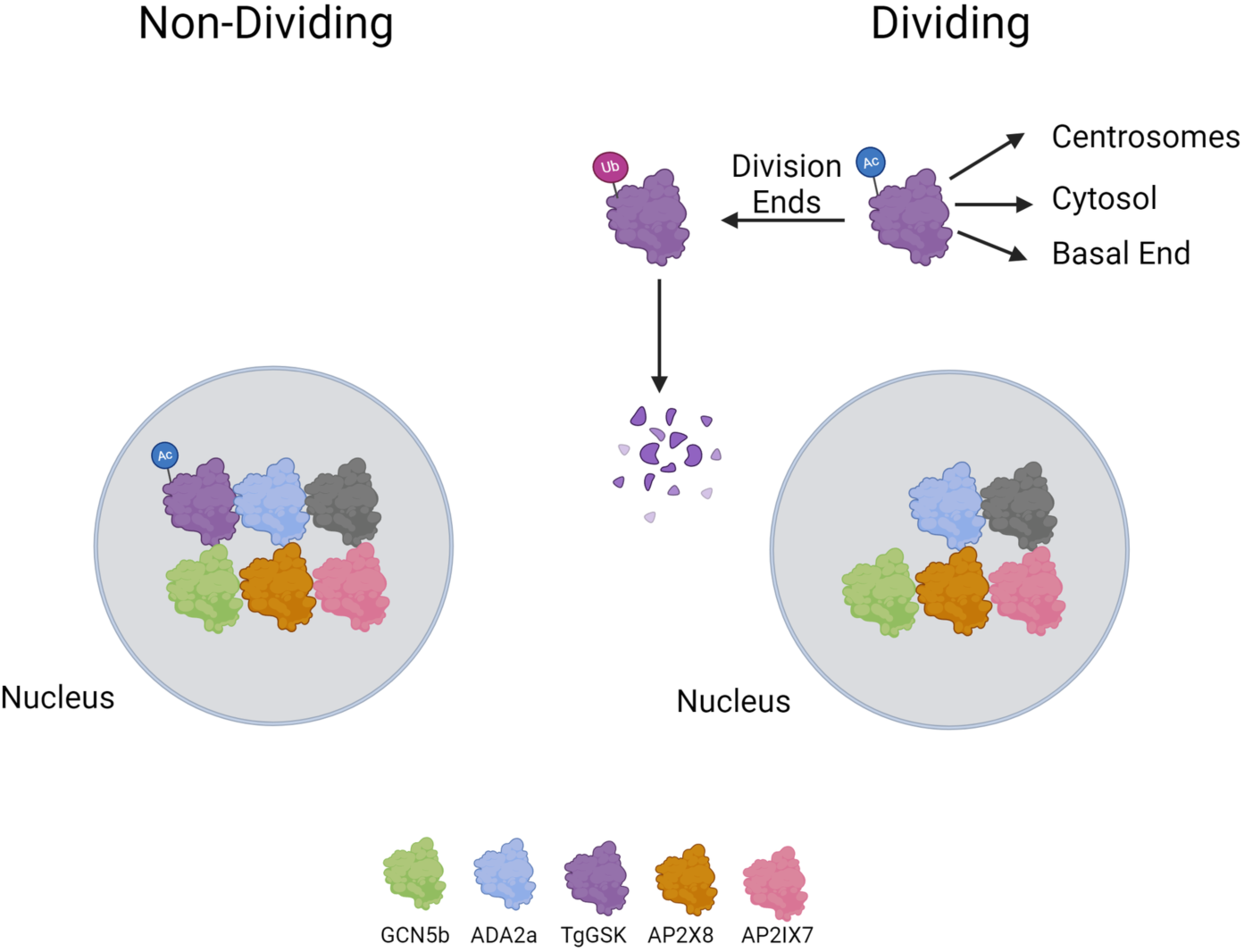
Model of the regulation and function of TgGSK. A preliminary model of TgGSK in non-dividing and dividing parasites. Image created using Biorender.

While phosphorylation in the activation loop is the best understood mechanism by which GSKs are regulated, a role for acetylation has also been reported. For example, mammalian GSK3β is acetylated at residue K183, and this acetylation is involved in kinase regulation [38]. A study of GSK3β in Alzheimer’s disease found that acetylation of GSK3β at residue K15 (the equivalent of TgGSK K13) led to the over-activation of the kinase, which led to the promotion of tau hyperphosphorylation and an increase in disease phenotypes [39]. In plants, HDAC6 removes acetylation on the GSK homolog BIN2 at residue K186 to inhibit kinase activity and enhance brassinosteroid signaling [40]. That study also showed that the acetylation and phosphorylation sites are both in the binding pocket and interact to regulate the kinase activity of BIN2 [40]. Interestingly, the residue shown to be acetylated in plants and mammals (K186 and K183, respectively) is conserved in *Toxoplasma*, but the acetylome data does not show that this residue is modified [41].

Overall, the findings in this study highlight the various roles of TgGSK in *Toxoplasma.* As BIN2 and mammalian GSKs have been shown to have hundreds of substrates with many different biological functions, the broad range of possible TgGSK functions uncovered here is not surprising. While likely roles for TgGSK in the centrosome and splicing have been identified through this work, further studies are warranted to understand the mechanistic underpinning of these functions. The critical importance of this essential kinase underscores its strong potential as a target for antiparasitic intervention.

## MATERIALS AND METHODS

### Parasite strains and reagents

All parasite strains used in this study derived from the strain RH lacking HXGPRT and Ku80 (RH*Δku80Δhxgprt*, referred to as *Δku80*) [42]. Parasites were maintained in human foreskin fibroblasts (HFF) with standard growth medium as previously described [43].

### Phylogenetic analysis

The GSKs used in the phylogenetic analysis include TgGSK (EPT27729), PfGSK (XP_001351197), AtBIN2 (Q39011), HsGSK3α (P49840), and HsGSK3β (P49841). Full protein sequence alignment and phylogenetic analysis were performed using Clustal Omega.

### Generation of parasite lines

All primers used for molecular cloning are listed in supplemental table 1. To add a hemagglutinin (HA) epitope tag to the endogenous TgGSK gene, we amplified the 3xHA-DHFR [44] amplicon from the LIC-3xHA-DHFR plasmid using primers that allowed for recombination at the 5’end with sequences immediately upstream of the stop codon and at the 3’end with sequences after the Cas9 cutting site. To direct these templates to the correct locus, we modified the plasmid pSag1-Cas9-U6-sgUPRT [45] using Q5 Site-Directed Mutagenesis Kit (NEB) to replace the UPRT guide RNA sequence within the plasmid to a guide RNA sequence of the target gene. The CRISPR/Cas9 plasmid and the PCR amplicon were transfected into parental parasites using the Lonza Nucleofector and the manufacturer’s suggested protocols. Transfected parasites were selected using pyrimethamine and cloned by limiting dilution as previously described [46].

To generate the TgGSK conditional knockdown strain, we utilized a CRISPR-Cas9 mediated strategy to introduce a tet-OFF cassette [47], which includes a transactivator (TATi) protein, the drug-selective marker HXGPRT, and a tet response element (TRE) followed by the Sag1 5’ UTR, immediately upstream of the TgGSK gene start codon. Specifically, a guide RNA targeting the TgGSK gene locus downstream of the start codon was constructed by mutating the plasmid pSag1-Cas9-pU6-sgUPRT [45] using the Q5 mutagenesis kit. The tet-OFF cassette was amplified from the vector pT8TATi-HXGPRT-tetO7S1 [47]. About 2 µg of the plasmid pSag1-Cas9-U6-sgGSK-KD and the PCR amplicon from 200 µl of PCR reactions with 30 cycles were transfected into the TgGSK-3xHA parasites using a Lonza nucleofector. Transfected parasites were then selected with 50 mg/mL mycophenolic acid (MPS) and xanthine and cloned by limiting dilution. Precise integration of the tet-OFF cassette was validated by PCR. The resulting strain was designated as TATi-GSK.3xHA. To induce knockdown of GSK, this strain was grown in 1µM of anhydrotetracycline (aTC) from Sigma Aldrich for the described length of time.

### Plaque assays

Standard plaque assays were performed as previously described [43]. Briefly, 500 parasites of each strain were seeded into host cell monolayers grown in 12-well plates, and cultures were then grown for six days. Cultures were then fixed with methanol and stained with crystal violet. Host cell plaques were quantified as previously described [43].

### Immunofluorescence assays

Immunofluorescence assays (IFAs) were performed as previously described [43]. The primary antibodies used include rabbit anti-HA (Cell Signaling Technologies), rat anti-IMC3 (provided by Dr. Marc-Jan Gubbels, Boston College), mouse anti-centrin 1 (Cell Signaling Technologies), and mouse anti-acetylated tubulin (Sigma Aldrich) at a concentration of 1:1000; guinea pig anti-TgEB1 (provided by Dr. Marc-Jan Gubbels, Boston College) at a concentration of 1:3000; rabbit anti-TgH2Bz (provided by Dr. Laura Vanagas and Dr. Sergio Angel, INTECH-Chascomus) at a concentration of 1:500; and rabbit anti-Cpn60 (provided by Dr. Erica Dos Santos Martins, UFMG) at a concentration of 1:300. Secondary antibodies used include Alexa Fluor 405, 488, 594, and 647 (Invitrogen) as well as DAPI (Thermo Fischer), all at 1:1000 or 1:2000. For images in figures 2B and C, 3B, 4A, and 9A, a Nikon Eclipse E100080i microscope with NIS Elements AR 3.0 software was used for imaging, followed by image analysis in ImageJ. For images in figures 4D, 5A, and 6A, a Zeiss LSM800 confocal microscope with Zeiss ZEN blue v2.0 and Huygens Professional v19.10.0p2 software was used for deconvolution, followed by image analysis in ImageJ.

Quantification of the GSK-HA signal in the nucleus and cytoplasm was performed by imaging 20 non-dividing vacuoles and 20 in late division using a Nikon Eclipse E100080i microscope with NIS Elements AR 3.0 software. Using ImageJ, fluorescent intensity was taken along a line in the cytosol and nucleus of a parasite in each vacuole before a ratio of nuclear to cytosolic fluorescent intensity was calculated for each vacuole. Figure 2D was made using GraphPad Prism software, and statistical analysis was performed using a student’s t-test.

### Ultrastructure expansion microscopy (UExM)

Parasites were grown on HFF monolayers on round coverslips and then fixed for 20 minutes in 4% paraformaldehyde. Coverslips were then treated with a 1.4% formaldehyde and 2% acrylamide solution in PBS overnight. Coverslips were inverted onto a solution of 5 µL 10% APS, 5 µL 10% TEMED, and 35 µL monomer solution (21% sodium acrylate solution, 28% acrylamide, 6% BIS, 11% 10x PBS) in a humid chamber and incubated on ice for five minutes and then at 37°C for one hour. Each gel and coverslip were then put in 2 mL of denaturation buffer (200 mM SDS, 200 mM NaCl, 50 mM Tris in water, pH 9) on the rocker for 15 minutes to remove the gel from the coverslip. Each gel was then placed in an Eppendorf tube with 1.5 mL of denaturation buffer and incubated at 95°C for 90 minutes. Each gel was then incubated three times in 25 mL of ddH_2_O for 30 minutes before being washed two times with 20 mL of 1x PBS for 15 minutes. Each gel was then blocked for 30 minutes in 3% BSA/PBS before incubation overnight with 1 mL primary antibody solution in 3% BSA/PBS at room temperature on the rocker. Gels were washed three times with 2 mL PBS-T for 10 minutes before incubation with 1 mL secondary antibody solution in PBS for 2.5 hours. Following another three PBS-T washes, gels were incubated again three times in 25 mL ddH2O for 30 minutes. Gels were then cut using the opened top of a 15 mL falcon tube and placed in 35mm Cellvis coverslip bottomed dishes that had been treated with poly-D-lysine. Primary antibodies used include rabbit anti-*Toxoplasma* tubulin (provided by Dr. Michael Reese, UT Southwestern), rabbit anti-centrin 1, and mouse anti-HA at a concentration of 1:500. DRAQ5 (1:500) and NHS Ester (1:250) were also used to visualize nuclear material and overall protein, respectively. Secondaries used include Alexa Flour 488 and 594 at a concentration of 1:500. Imaging was performed using a Zeiss LSM900 microscope with Zeiss ZEN Blue software before image analysis using ImageJ.

### Western blots

Western blots were performed as described previously [43]. The primary antibodies used include rabbit anti-HA (Cell Signaling), mouse anti-Sag1 (Invitrogen), and mouse anti-aldolase. The secondary antibodies utilized were HRP-labeled Anti-Mouse and Anti-Rabbit IgG. The primary antibodies were used at a dilution of 1:5,000, while the secondary antibodies were used at a dilution of 1:10,000. Imaging of the blot was performed using a ProteinSimple system.

### Immunoprecipitation

Immunoprecipitation was performed as previously described with some modifications [46]. For immunoprecipitation from whole-parasite lysate, intracellular parasites of the GSK.3xHA and Ku80 strains were grown for 18 hours in host cells. Parasites were harvested with host cells by scraping in cold PBS and centrifugation at 2000 rcf for 5 minutes at 4°C. Cells were lysed with 500 µL RIPA lysis buffer supplemented with 5µL protease and phosphatase inhibitor cocktail (Thermo Scientific) at 4°C for one hour. Each sample was sonicated three times and centrifuged at maximum speed for 10 minutes at 4°C. The supernatant of each sample was incubated with mouse IgG magnetic beads for one hour at 4°C for pre-cleaning. Samples were then incubated with rabbit HA magnetic beads (Thermo Scientific) overnight at 4°C. After washing with RIPA lysis buffer and PBS, the beads were submitted to the Indiana University School of Medicine Proteomics Core facility for liquid chromatography coupled to tandem mass spectrometry (LC/MS-MS) analysis.

### Global transcriptomic analysis

TATi-GSK.3xHA parasites were grown for 18 hours with or without aTC in host cells. Parasites were harvested with host cells by scraping in cold PBS, followed by centrifugation at 2000 rcf for five minutes at 4°C. The pellet was passed through a syringe in 10 mL PBS to release parasites from host cells, and the samples were centrifuged again. The pellet was treated with 1 mL TRIZOL for five minutes at room temperature before extracting RNA with 200 µL of chloroform and centrifuging at 12,000 rcf for 15 minutes at 4°C. The aqueous phase was again treated with 500 µL of chloroform and centrifuged to extract as much RNA as possible. The aqueous phase was then mixed with 500 µL of isopropanol and incubated at room temperature for 10 minutes before centrifuging at 12,000 rcf for 10 minutes. The RNA was washed with 1 mL 75% ethanol and again centrifuged at 7,500 rcf for five minutes. The resulting RNA pellet was dried and resuspended in 50 µL of nuclease-free water. Each condition was performed in triplicate. Samples were stored at -80°C before being sent to AZENTA for library construction and sequencing utilizing Illumina Next Generation Sequencing technology. For each sample, ∼30 M 2x150 bp pair-end reads were obtained. The GALAXY online platform was used to perform data analysis. Specifically, the quality of the sequencing data was checked using FastQC, and adapter sequences were trimmed using Trim Galore. Hisat2 and htseq-count were separately employed to map reads to the genome and count the reads of each transcript. DEseq2 was used to analyze differential gene expression, and DEXseq was utilized to analyze whether the knockdown of TgGSK affects alternative splicing. Pathway analysis was done through a combination of ToxoDB and StringDB.

### Global phosphoproteomics

TATi-GSK.3xHA parasites were grown for 24 hours with or without aTC in host cells. Parasites were harvested via scraping in cold PBS and centrifuged at 2000 rcf for five minutes at 4°C. Parasites were released from host cells using a syringe with a 27-gauge needle, and the samples were centrifuged again. Each condition was performed in triplicate. The parasite pellets were flash-frozen in liquid nitrogen and stored at -80°C before being sent to the Indiana University School of Medicine Proteomics Core for global phosphoproteome analysis, and they performed sample preparation and analysis as described previously [6]. Protein function identification and pathway information were determined using ToxoDB and StringDB.

### Garcinol assays

GSK.3xHA parasites were seeded simultaneously with 2 or 4 µM Garcinol (BOC Sciences) in 1% serum media for 18 hours. Immunofluorescence analysis and western blots were done as described above.

## ACKNOWLEDGMENTS

The mass spectrometry work was performed by the Indiana University School of Medicine Center for Proteome Analysis. This research was supported by the National Institutes of Health grants R01AI149766, R01DK124067, and R21AI164619 to G.A. The Lab. of Apic. Biol. at Institut Pasteur de Montevideo is funded by a G4 grant awarded to MEF by the Pasteur Network and FOCEM (MERCOSUR 645 Structural Convergence Fund), COF 03/11. The authors gratefully acknowledge the Advanced Bioimaging Unit at the Institut Pasteur Montevideo. MEF is a SNI and PEDECIBA researcher.

---

**Supplemental table 1.**
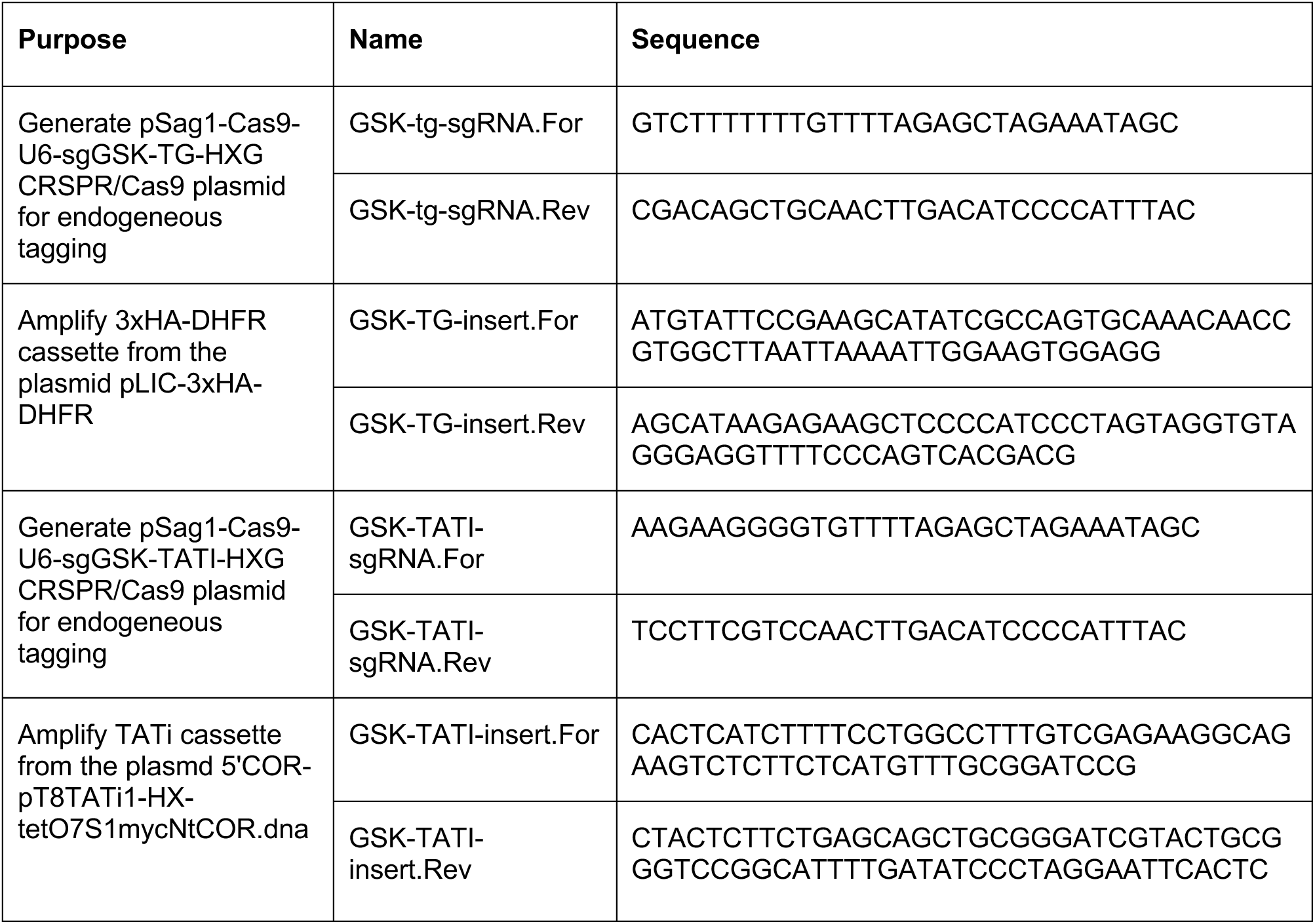
Primers used in this work. Sequences are 5’ to 3’.

**Supplemental figure S1.**
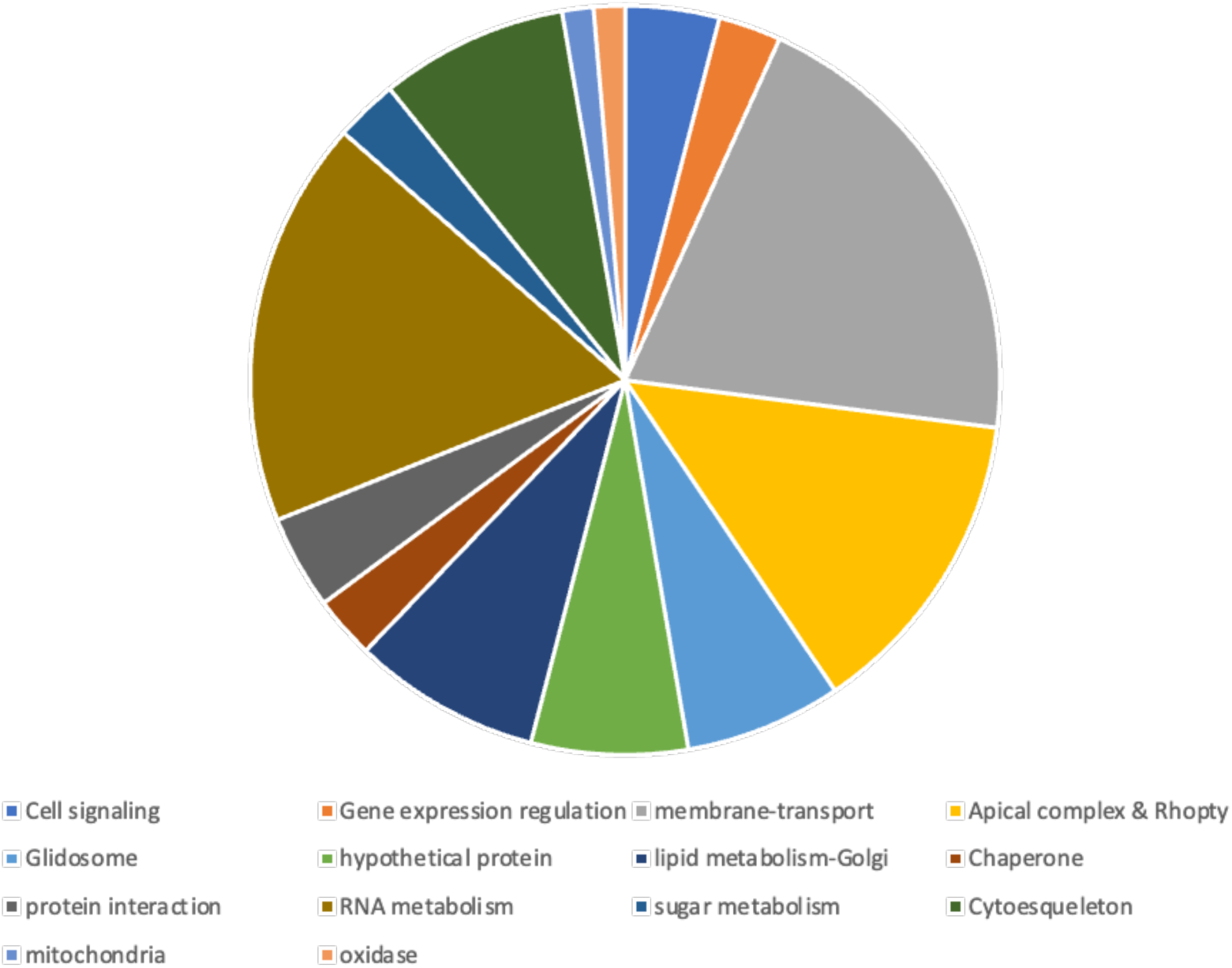
Classification of hypothetical proteins with TgGSK-dependent phosphopeptides. Hypothetical proteins with peptides differentially phosphorylated in the knockdown vs the parental were analyzed based on annotations in the *Toxoplasma* genome database or homology to proteins in other Apicomplexan species. Additionally, the protein domains were analyzed based on known conserved functions. Some of these proteins remained classified as hypothetical.

## Notes

### Competing Interest Statement

The authors have declared no competing interest.

